# Lumen expansion facilitates epiblast-primitive endoderm fate specification in the mouse blastocyst formation

**DOI:** 10.1101/575282

**Authors:** Allyson Quinn Ryan, Chii Jou Chan, François Graner, Takashi Hiiragi

## Abstract

Mouse blastocyst formation involves lumen formation and cell fate specification. While many studies have investigated how the blastocyst cell lineages are specified through genetics and signaling, studies into the potential role of the fluid lumen have yet to be conducted. We discover that blastocyst fluid emerges by secretion of cytoplasmic vesicles to intercellular space in addition to trans-epithelial flow. We observe that the beginning of epiblast and primitive endoderm spatial segregation directly follows lumen coalescence. Notably, we show that perturbing lumen expansion by pharmacological and biophysical means impair the specification and spatial segregation of primitive endoderm cells within the blastocyst. Combined, our results suggest that blastocyst lumen expansion plays a critical role in guiding cell fate specification and positioning. As epithelial tissues typically form lumina, lumen expansion may provide a general mechanism of cell fate control in many tissues.

## Introduction

Pre-implantation mouse development culminates with the formation of a blastocyst containing three spatially segregated cell lineages and an abembryonically localized fluid lumen. The molecular specification of the three cell lineages – trophectoderm (TE), epiblast (EPI) and primitive endoderm (PrE) – occurs sequentially (Artus et al., 2011; Chazaud et al., 2006; Rossant and Tam, 2009). Asymmetric divisions and differential contractility result in a 16-cell stage embryo containing apolar inner that upregulate inner cell mass (ICM) markers and polar outer cells that upregulate TE markers (Anani et al., 2014; Johnson and Ziomek, 1981; Korotkevich et al., 2017; Maître et al., 2016; Nishioka et al., 2009; Strumpf et al., 2005). Subsequent cleavage rounds to 32, 64 and 128-cell stages see the establishment of the epiblast (EPI) and primitive endoderm (PrE) cell lineages. During the 32-cell stage, inner cells are stochastically biased toward either EPI or PrE identity on a molecular level (Chazaud et al., 2006; Guo et al., 2010; Kang et al., 2013; Ohnishi et al., 2014; Schrode et al., 2014). These biases are either reinforced or changed during the remainder of preimplantation development depending on cell rearrangements and final cell position within the ICM, such that the EPI cells are situated between TE and PrE cells, which align along the luminal surface (Frankenberg et al., 2011; Plusa et al., 2008; Saiz et al., 2013).

Concurrently with the specification of EPI and PrE, the blastocyst lumen begins to form and expand (Rossant and Tam, 2009). The major driver of fluid accumulation within the embryo is thought to be Atp1, a Na^+^/K^+^-ATPase, which forms an osmotic gradient across TE cells through its polarized basolateral expression (Wiley, 1984; Watson, 1992; Watson and Barcroft, 2001). Early phases of fluid accumulation result in multiple fluid pockets that preferentially localize at the base of TE cells before coalescing to form the abembryonic pole of the embryonic-abembryonic axis (Figure 1A; Motosugi et al., 2005). The apical domain of TE cells continues to face the external environment of the embryo throughout lumen expansion, which is inverted to the morphology of a typical cyst where the apical side is facing the lumen (Bryant and Mostov, 2008). Inner cells exposed to the lumen do not undergo polarization and epithelialization until the very end of preimplantation development (E3.75-E4.0; Gerbe et al., 2008). Tight regulation and maintenance of the established apico-basal polarity and tight junctions within the TE cells are required for blastocyst lumen formation and expansion regulation (Eckert et al., 2004; Madan et al., 2007; Moriwaki et al., 2007).

**Figure 1.**
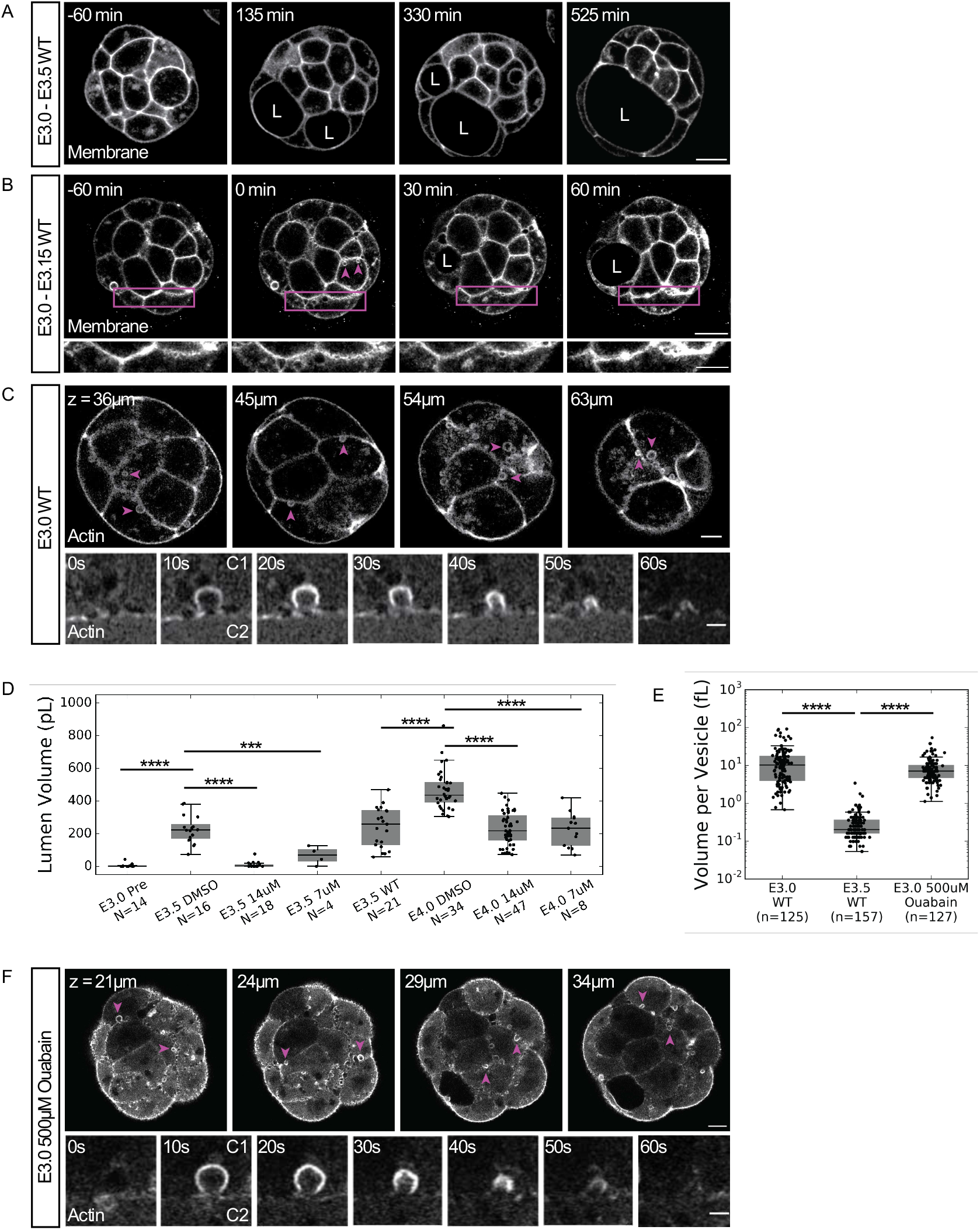
Blastocyst cavities are partially derived from cytoplasmic vesicles. (A) Time-lapse of a representative embryo expressing a membrane marker undergoing lumen formation (L marks a lumen). *t* = 0min when fluid accumulation is first detectable by automatic segmentation. Scale bar = 20μm. (B) Time-lapse of the first hour of fluid accumulation in an embryo expressing a membrane marker (L marks a lumen). *t* = 0min when ‘string of pearls’ microluminal structures are observed. Top row is full embryo view (magenta arrowheads highlight cytoplasmic vesicles, scale bar = 20μm). Bottom row is insets of cell-cell interfaces indicated by magenta boxes in top row highlighting the appearance of ‘string of pearls’-like microlumina emergence and resolution (scale bar = 10μm). (C) Z-slices of phalloidin staining showing cortically-localized vesicles in an E3.0 embryo (top, magenta arrowheads highlight individual vesicles, scale bar = 10 μm). Time-lapse of vesicle secretion into intercellular space in a Lifeact-GFP E3.0 embryo (bottom, ‘C1’ marks secreting cell, ‘C2’ marks adjacent cell, scale bar = 2μm). (D) Box plot of volume for lumina in Brefeldin A pharmacologically inhibited embryos. (E) Box plot of volume for individual vesicles in WT embryos at E3.0 and E3.5, and ATP1 inhibited embryos at E3.0. (F) Z-slices of an E3.0 embryo expressing Lifeact-GFP showing vesicle localization under Atp1 inhibition conditions (top, magenta arrowheads highlight individual vesicles, scale bar = 10μm). Time-lapse of vesicle secretion into intercellular space under Atp1 inhibition conditions (bottom, ‘C1’ marks secreting cell, ‘C2’ marks adjacent cell, scale bar = 2μm). ***p<0.001 ****p<0.0001. N = embryo N = vesicle

Thus far, the function(s) of the blastocyst lumen are unknown. Depending on the tissue and developmental time, lumina have widely varied functions including cell shape change (Gebala et al., 2016), nutrient delivery and absorption (Sobajima et al., 2015), left-right symmetry breaking (Essner et al., 2005) and even signaling niche establishment (Durdu et al., 2014). Interestingly, it has been shown in the mouse pancreas that failed tubulogenesis causes altered fate allocation ratios of progenitor cells due to changes in microenvironments (Kesavan et al., 2009). Despite the temporal correlation of EPI-PrE specification and lumen formation, the potential function of the lumen to regulate fate specification and cell positioning have yet to be investigated. In this work, we examined the early phases of blastocyst lumen formation and expansion in relation to PrE cell initial specification and migration with an aim to better understand the interplay between cell fate specification and blastocyst morphogenesis.

## Results

### Widespread secretion of cytoplasmic vesicles into intercellular space drives early fluid accumulation

Given the multipoint origin of the blastocyst lumen (Motosugi et al., 2005), we examined the first moment of extracellular fluid accumulation at high spatial resolution over multiple timescales (Figure 1A-C). In addition to the multiluminal stage, we observed the unbiased appearance of microlumina arranged in a ‘string of pearls’ like morphology at cell-cell interfaces throughout the embryo that progressively undergo coalescence over 2-3 hours (Figure 1B). In embryos at a stage just prior to measurable separation of cell membranes due to fluid accumulation (approximately 84hrs post-hCG, E3.0), we observed large, cortically-localized, actin-coated vesicles along the basolateral membranes of outer cells and ubiquitously along the membranes of apolar, inner cells (Figure 1C, top panel; Movie S1). These vesicles are actively secreted into intercellular space in approximately 60 seconds (Figure 1C, bottom panel; Movie S2). The presence and dynamic behavior of these vesicles persists through the early phases of luminal coalescence and expansion (E3.0-E3.25); however, such vesicles are no longer observable in embryos in which the lumen has expanded to occupy at least 50% of the total embryo volume (approximately 96hrs post-hCG, E3.5; Figure 1E, Figure S1). Actin coated vesicles in E3.5 embryos are significantly smaller than those in E3.0 embryos; they form cytoplasmic clusters, and no observable secretion events occur at the same time scale as that of E3.0 vesicles (Figure 1 E, S1).

To discern if vesicle release makes a measurable contribution to total luminal volume, embryos from early (E3.0-E3.5) and late (E3.5-E4.0) stages of lumen expansion were incubated in media containing Brefeldin A, a well-known inhibitor of COPII machinery (Miller et al., 1992; Helms and Rothman, 1992) that does not affect cell divisions (Figure S2). Titrations of the Brefeldin A revealed that inhibitory effects on luminal volume can be modulated during early lumen expansion phases (Figure 1D) in agreement with the observation of vesicle release occurring primarily during early expansion (Figure 1C, Figure S1). To determine if vesicle release is linked to the Atp1 driven mechanism of fluid accumulation, we incubated embryos with ouabain, a well-known inhibitor of Atp1 activity (Hoijman et al., 2015; Manejwala et al., 1989), for two hours prior to fluid accumulation onset. We observed that the vesicle release mechanism is still present in embryos cultured with ouabain (Figure 1F, top panel; Figure 1E; Movie S3). The time it takes a single vesicle in Atp1 inhibited embryos to be secreted appears similar to that of WT embryos (Figure 1F, bottom panel; Movie S4). These results indicate that vesicle secretion as a fluid accumulation mechanism is independent of Atp1.

### Early luminal structures are marked with apical proteins and contain FGF4

While the apicobasal polarity of TE cells surrounding the mouse blastocyst lumen is inverted to that of typical cysts and tubes (Alvers et al., 2014; Bryant et al., 2010 and 2014), the vesicle fusion observed in E3.0 embryos is similar to exocytosis of apical vacuolar compartments in apical cord hollowing (ACH), which is a *de novo* lumen formation mechanism that is conserved across species and tissues (Alvers et al., 2014; Bryant and Mostov, 2008; Sigurbjörnsdóttir et al., 2014). Critical to the initiation of ACH is the formation of the apical membrane initiation site (AMIS) which dictates where the lumen will form and expand (Bryant et al., 2010; Ferrari et al., 2008) through exocytosis of apical vacuolar compartments (Bryant and Mostov, 2008). As such, we examined early lumen formation stage embryos for apical polarity phenotypes resembling reported AMIS and AMIS-like structures. Interestingly, we found that many E3.0 embryos contain microlumina enriched for the apical marker phosphorylated ERM (pERM) (43%, N = 20 of 47 embryos, Figure 2A,B). By E3.25 (90hrs post-hCG), such structures are rare as the main lumen expands and individual microlumina merge with it (Figure 2B).

**Figure 2.**
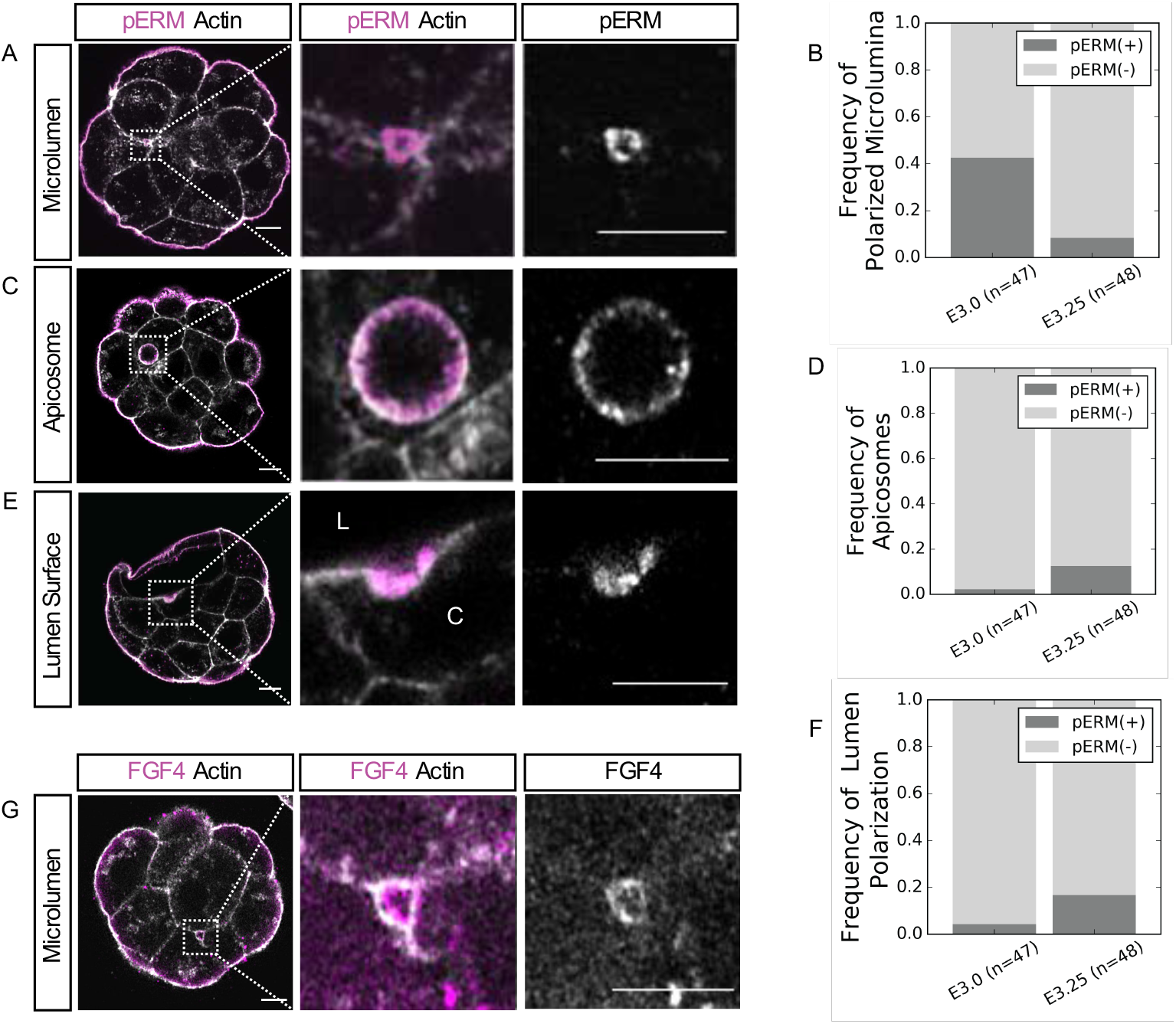
Microlumina containing secreted apical domain components are transiently upregulated during early phases of fluid accumulation. (A) Representative immunofluorescence images of an apically polarized microlumina in an E3.0 embryo. (B) Frequency of apically polarized microlumina in E3.0 and E3.25 embryos (p <0.001). (C) Representative immunofluorescence image of an E3.25 ICM cell containing an apicosome. (D) Frequency of apicosome occurrence in E3.0 and E3.25 embryos (p < 0.115). (E) Representative immunofluorescence image of an E3.25 ICM cell in which a subsection of its membrane facing the growing lumen is apically polarized (L – lumen; C – cytoplasm). (F) Frequency of lumen polarization in E3.0 and E3.25 embryos (p <0.091). (G) Z-slice of an RNA-injected E3.0 embryo showing localization of FGF4-mNeonGreen in a microlumen, representative of N = 7 embryos. All scale bars = 10μm. Two-tailed Fisher exact test.

While apically polarized microlumina are infrequent in E3.25 embryos, we do observe the presence of two other apically polarized structures rarely found in E3.0 embryos (Figure 2C-F). A small number of ICM cells in E3.25 embryos contain apicosome-like structures (13%, N = 6 of 48 embryos, Figure 2C,D). Apicosomes are structures containing apical polarity proteins described in human pluripotent stem cells, and have been proposed to be luminal precursors (Taniguchi et al., 2017). Cells containing apicosome-like structures are isolated from any contact-free surfaces created by the growing lumen (Figure 2C, Movie S5). If such cells acquire sustained contact with the lumen the apicosome is released into the lumen over approximately 2-3 hours postcontact (Movie S6). Subsections of the lumen-facing membrane in a small number of E3.25 ICM cells express apical polarity markers and have a markedly shorter radius of curvature than that of the rest of the ICM-lumen interface indicating recent fusion of either an apicosome or apically polarized microlumina (17%, N = 8 of 48 embryos, Figure 2E).

FGF4 has been shown to be essential for PrE establishment (Kang et al., 2013; Krawchuk et al., 2013; Lanner and Rossant, 2010; Yamanaka et al., 2010), and its expression is restricted restricted to EPI cells (E3.25-E4.5) (Frankenberg et al., 2011; Ohnishi et al., 2013). Because FGF ligands have been shown to create signaling niches by localizing to microlumina (Durdu et al., 2014), we examined the localization of FGF4 protein in E3.0 embryos by injecting mRNA of *fgf4-mNeonGreen* into a single blastomere of a 4-cell stage embryo. We observe localization on the membranes of a subset of microlumina (36%, N = 7 embryos, Figure 2G). The localization of FGF4 to microlumina suggests the possibility of luminal microenvironments capable of impacting fate specification in the surrounding cells.

### ICM spatial patterning resolves as the lumen expands

During the final phases of coalescence and lumen expansion, initial transcriptional biases of ICM cells towards either EPI or PrE fate are either reinforced or changed concomitantly with spatial segregation (Chazaud et al., 2006; Frankenberg et al., 2011; Ohnishi et al., 2014). Cells that are molecularly specified to become PrE but fail to achieve correct positioning along the ICM-lumen interface in a timely manner undergo apoptosis (Plusa et al., 2008). However, it is not known precisely when the cells begin to undergo repositioning since initial position of the precursors is stochastic (Chazaud et al., 2006; Gerbe et al., 2008), and lumen formation onset is variable in terms of absolute developmental time. To determine whether spatial segregation is correlated with luminal volume, we used lineage reporters and membrane signal (Muzumdar et al., 2007) to track the emergence and positioning of EPI precursor cells (Arnold et al., 2011) and PrE precursor cells (Hamilton et al., 2003) within the ICM in relation to the expanding lumen.

In order to measure and track the EPI and PrE domains accurately, we first developed an analysis method that incorporates gross morphological changes occurring over the course of lumen formation and expansion. The resulting method allows us to determine the distance of each ICM lineage from the ICM-lumen interface relative to the volume of the expanding lumen and the changing morphology of the ICM by simultaneously segmenting the lumen and embryo cell mass and determining the embryonic-abembryonic axis to provide spatial orientation (Figure 3A; see Methods for details).

**Figure 3.**
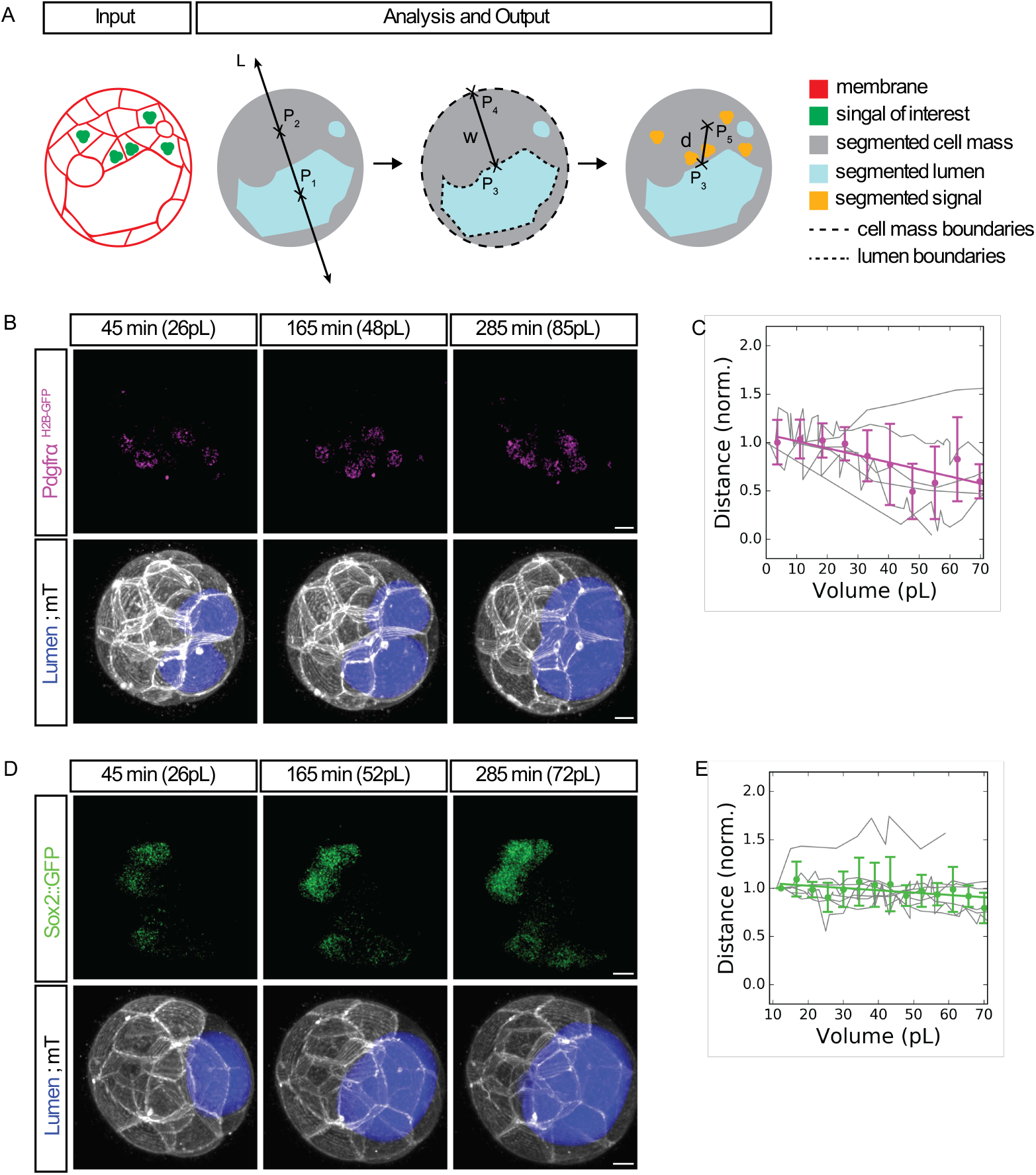
The Pdgfrα signaling domain approaches the luminal surface as the lumen grows in volume. (A) Schematic 2D representation of 3D analysis method for the tracking and normalization of fate reporter expression proximity to the ICM-lumen interface. *P*_1,2,3,4,5_ are 3D points. 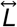 is a 3D line (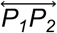 equivalent) that defines the embryonic-abembryonic axis. 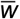 is the 3D line segment (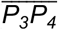 equivalent) that measures the ICM width. 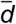 is the 3D line segment (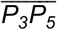 equivalent) that measures the distance from the center of mass of the signal of interest to the ICM-lumen interface. See Image Analysis for formal definitions of all geometric entities. (B) Time-lapse of an E3.0 embryo expressing a PrE reporter (*Pdgfrα^H2B-GFP/+^*; top), a membrane marker and lumen segmentation (bottom). *t* = 0min is defined as the first moment when a lumen can be segmented. Lumen volume (pL) is given for each time point shown. (C) Quantification of the distance of the center of Pdgfrα signaling domain to the surface of the lumen over time (N = 6 embryos, thin gray lines are traces of individual embryos, magenta dots are binned averages with vertical capped lines showing standard deviations, thick magenta line is the linear regression y = −0.007x + 1.090, r^2^ = 0.653, p < 0.005). (D) Time-lapse of an E3.0 embryo expressing a cytoplasmic EPI reporter *(Sox2::gfp*; top), a membrane marker and lumen segmentation (bottom). *t0* is defined as the first moment when a lumen can be segmented. Lumen volume (pL) is given for each time point shown. (E) Quantification of the distance of the center of Sox2 expression domain to the surface of the lumen over time (N = 8 embryos, thin gray lines are traces of individual embryos, green dots are binned averages with vertical capped lines showing standard deviations, thick green line is the linear regression y = −0.002x + 1.074, r^2^ = 0.339, p < 0.030). All scale bars = 10μm.

The EPI reporter (Sox2::gfp) center showed no net movement toward the ICM-lumen interface and maintained a relative constant distance from the luminal surface throughout expansion (Figure 3D,E). In contrast, the PrE reporter (Pdgfrα^H2B-GFP/+^) center shifts toward the ICM-lumen interface as soon as coalescence begins (approximately E3.25) and continues throughout expansion (Figure 3B,C). This indicates differential sorting behavior between PrE and EPI cells, and suggests that lumen expansion may play a role in guiding EPI-PrE fate specification and spatial segregation.

### ICM lineage specification and spatial segregation are dependent on luminal expansion

Given the correlation between lumen expansion with the specification and subsequent spatial segregation of EPI and PrE progenitor cells, we hypothesized that modulation of lumen size may impair lineage differentiation and cell positioning. To test this hypothesis, we inhibited the expansion of post-coalescence stage lumina (96-108 hours post-hCG; E3.5-E4.0) through multiple means. Embryos cultured in media containing 500μM ouabain (Wiley, 1984) show significant decrease in luminal volume. The volume effect is titratable by changing inhibitor concentration (Figure 4A,B; Figure S3A), suggesting the effect of ouabain in reducing lumen expansion is specifically due to its action on the Atp1 channel. Notably, these embryos show significant reduction in the expression levels of both EPI and PrE markers in addition to possessing a significantly lower luminal volume than that of controls (Figure 4) without a change in the total number of cells (Figure S3B).

**Figure 4.**
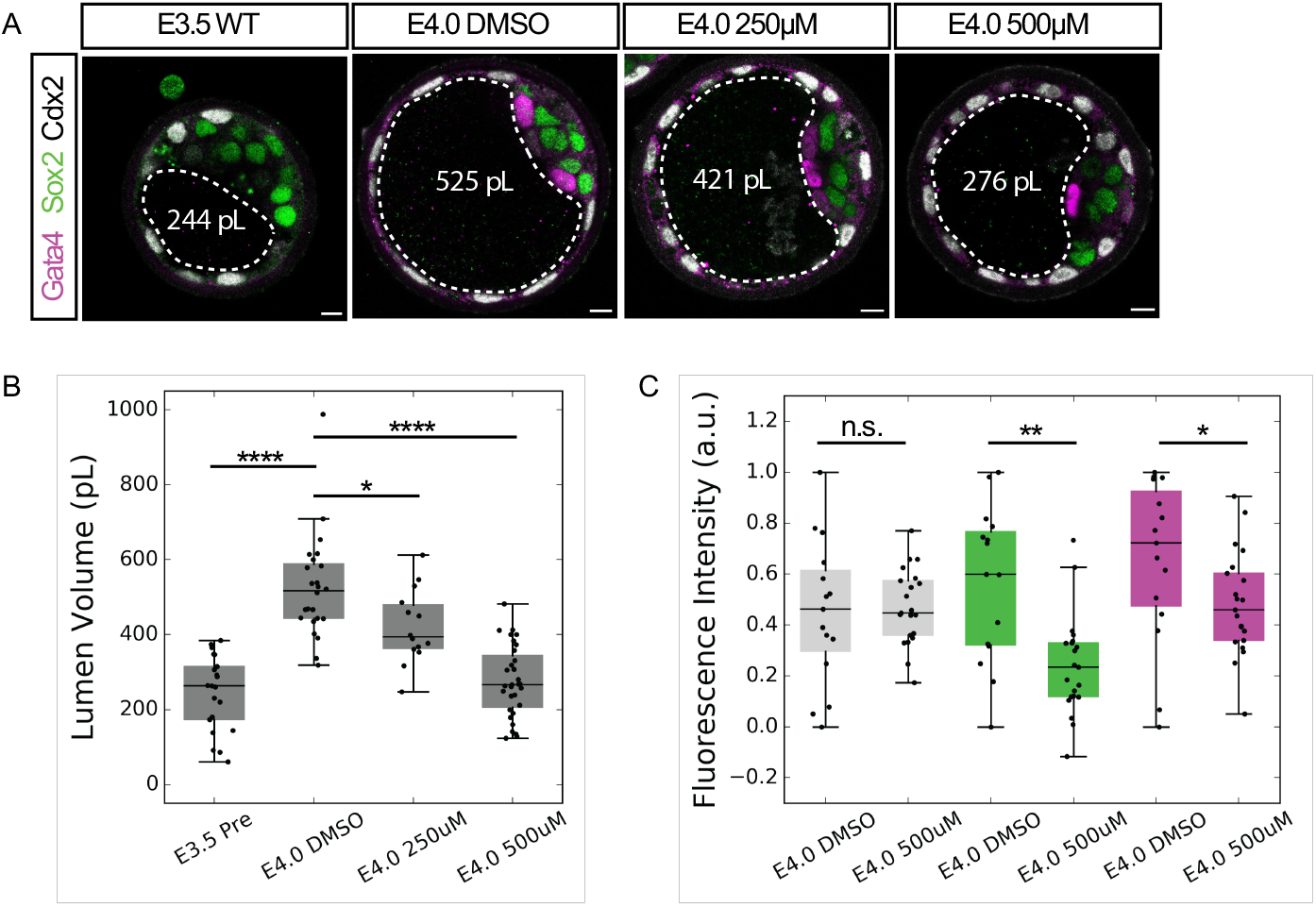
EPI and PrE expression levels are reduced in ATP1 inhibited embryos. (A) Immunofluorescence images of TE (Cdx2), EPI (Sox2) and PrE (Gata4) fate in pre-treatment control (E3.5 WT), Atp1 inhibited (E4.0 500μM and E4.0 250μM), and end-stage control (E4.0 DMSO) embryos. Lumen boundaries outlined by dashed white line and mean lumen volume in white text. Scale bars = 10μm. (B) Boxplot of lumen volume for E3.5 WT (N = 21), E4.0 DMSO (N = 24), E4.0 250μM Atp1 inhibited (N = 14) and E4.0 500μM Atp1 inhibited (N = 31) embryos indicating that the impact on lumen volume is concentration dependent. (C) Boxplot of fluorescence levels of Cdx2 (gray), Sox2 (green) and Gata4 (magenta) in E4.0 500μM Atp1 inhibited embryos compared to E4.0 dMsO controls. ****p<0.0001 **p<0.01 *p<0.05.

To further examine the possible role of lumen expansion in cell fate specification and sorting, we mechanically deflated late expansion stage lumina (E3.5-E4.0) by inserting a microneedle into the lumen at the junctions of mural TE cells, and applying negative pressure to block expansion (Figure 5A). This action was repeated every 1-2 hours as necessary for individual embryos so that the blastocyst lumen volume did not exceed initial E3.5 volumes. To ensure observations are not due to adverse effects induced by serial needle insertion, we performed a control of passive lumen deflation due to serial puncture (‘Control’, Figure 5E). Importantly, serial needle insertion does not perturb embryo cell number (Figure S4).

**Figure 5.**
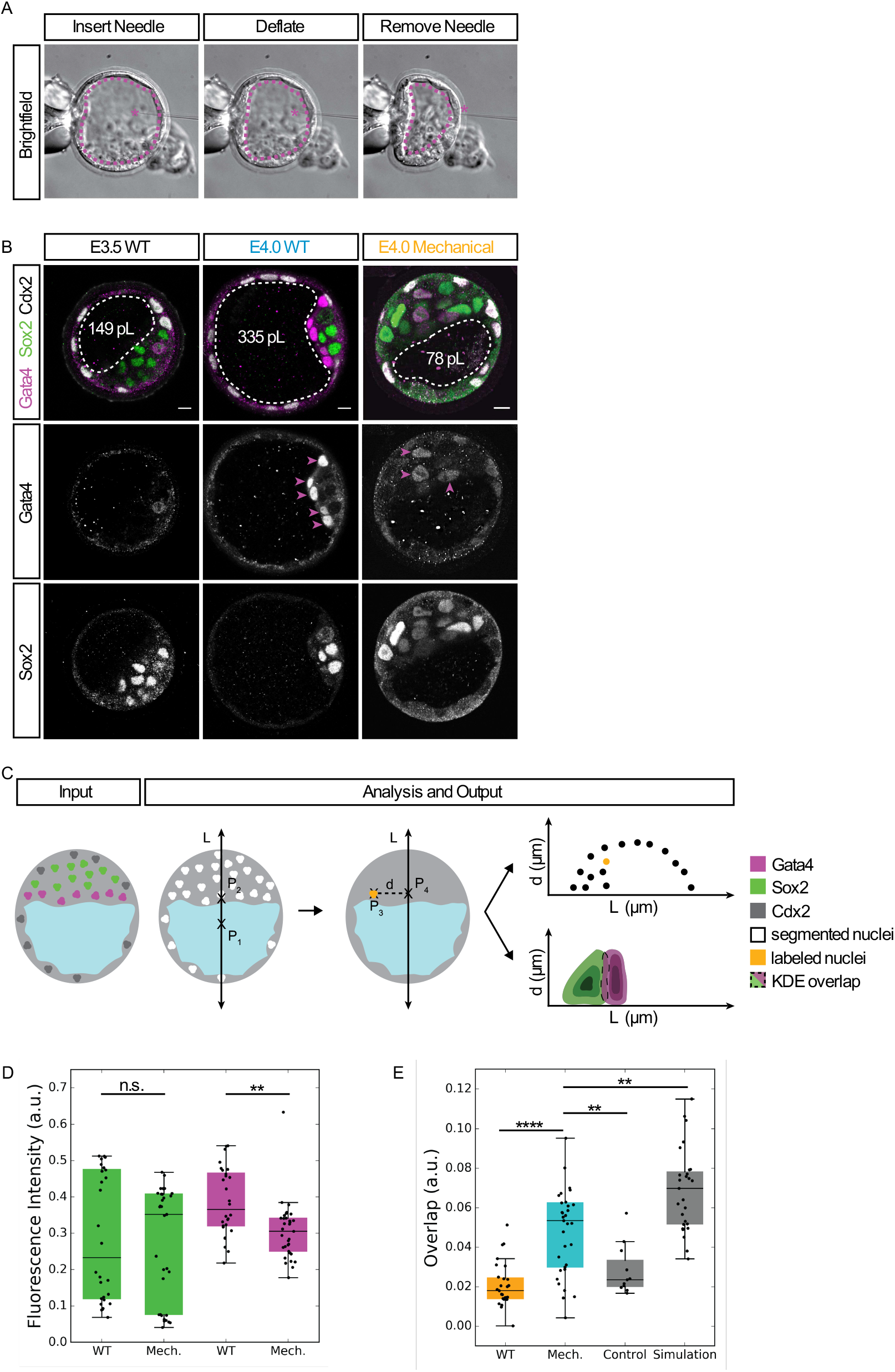
PrE specification and spatial segregation of ICM lineages is impaired by mechanical inhibition of lumen expansion. (A) Brightfield images of mechanical deflation. Magenta asterisk marks the needle tip. Dotted magenta line indicates lumen boundary. (B) Immunofluorescence images of EPI (Sox2) and PrE (Gata4) fate in pre-manipulation control (E3.5 WT), E4.0 post-manipulation control (E4.0 WT) embryos and E4.0 mechanically inhibited (E4.0 Mechanical). Magenta arrowheads indicate the position of cells expressing high levels of Gata4 within the ICM. White dotted line indicates lumen boundaries. Average lumen volume in white text. Scale bars = 10μm. (C) Schematic 2D representation of 3D analysis method for spatial segregation of ICM lineages. *P*_1,2,3,4_ are 3D points. 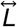 is a 3D line (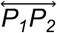 equivalent) that defines the embryonic-abembryonic axis. 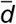 is the 3D line segment (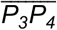 equivalent) that measures the perpendicular distance from the center of a cell to 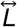. See Image Analysis for formal definitions of all geometric entities. (D) Boxplot of fluorescence levels of Sox2 (green) and Gata4 (magenta) in mechanically inhibited (‘Mech’, N = 33) and post-manipulation control (‘WT’, N = 28) E4.0 embryos. (E) Boxplot of spatial overlap between EPI and PrE lineages within post-manipulation control (‘WT’, N = 27), mechanically inhibited (‘Mech’, N = 33), E4.0 procedural control (‘Control’, N = 11) and E4.0 simulation of complete overlap in WT conditions (‘Simulation.’, N = 27). ****p<0.0001 **p<0.01 n.s. = not significant

Interestingly, PrE specification levels are significantly reduced by mechanical deflation, while EPI specification levels are maintained at the level of WT embryos (Figure 5B,D). Additionally, mechanical deflation impairs the spatial segregation of EPI and PrE cells (Figure 5B,E). EPI-PrE overlap is defined as the intersection of the probability of the two populations based on spatial position in 3D and expression level of Sox2 (EPI) and Gata4 (PrE) (Figure 5C; see Methods for details). EPI-PrE overlap of mechanically deflated embryos is more than doubled in comparison to WT controls (mean_Mech_= 0.053, mean_WT_ = 0.021, p<0.0001), but is still significantly less than that of simulated maximal overlap values (‘Simulation’, mean_Simulation_ = 0.069, p < 0.005). Maximum potential overlap (‘Simulation’) is defined as the scenario in which the entire ICM is equally likely to be EPI or PrE, resulting in a high overlap value. Spatial segregation analysis of controls shows a slightly higher degree of overlap between EPI and PrE domains in comparison to that of WT (mean_Control_ = 0.028, p < 0.045) while still being significantly lower than that of mechanically deflated embryos (p < 0.007).

Combined, the results of pharmacological and mechanical inhibition consistently show that timely lumen expansion facilitates fate specification and spatial segregation of EPI-PrE cell lineages.

## Discussion

Collectively, the data presented here reveal a new mechanism of blastocyst lumen formation and its role in facilitating the specification and positioning of the PrE. We have shown that all cells of the embryo, not only the trophectoderm, can contribute to lumen initiation and growth through a vesicle secretion mechanism (Figure 1), in agreement with previous notions (Wiley and Eglitis, 1980 and 1981; Fleming and Pickering, 1985; Aziz and Alexandre, 1991). Furthermore, we have shown that immediately following sufficient coalescence of microlumina into a singular lumen, PrE biased ICM cells begin directional movement toward the ICM-lumen interface (Figure 3). When lumen expansion is perturbed by various methods, PrE maturation and repositioning is disrupted (Figure 4,5), suggesting a causal role of lumen expansion in guiding EPI-PrE cell fate specification and spatial sorting. Mechanistically, the presence of secreted molecules, effecting both polarity and fate specification, may suggest an instructive role of the luminal fluid contents (Figure 2).

Segregation of PrE-cells to delineate the spatial domains of EPI and PrE is crucial for tissue organization in downstream embryonic development (Moore et al., 2014; Morris et al., 2002; Yang et al., 2007). Here we observe oriented movement of the primitive endoderm toward the ICM-lumen interface immediately following luminal coalescence (E3.25, Figure 3). This result indicates that spatial segregation of ICM cell lineages initiates earlier than previously thought and depends crucially on morphological changes within the embryo. The oriented movement of primitive endoderm cells in accordance with lumen expansion (Figure 3–5) is reminiscent of chemokine-guided migration (Boldajipour et al., 2008; Bussman and Raz, 2015; Dona et al., 2013). Key to directed migration is the presence of either a biochemical or physical cue to provide orientation. Net movement of PrE primed cells but not EPI primed cells suggests differential reception of luminal cues by ICM cells. In this study we reported the presence of FGF4 in early luminal structures (Figure 2). It will be interesting for future studies to investigate the role of these signaling molecules as potential chemoattractants for PrE cells.

The presence of apically polarized microlumina in E3.0 embryos (Figure 2A) along with the widespread vesicle release (Figure 1C) reveal notable parallels between blastocyst lumen formation mechanisms and ACH. The apicosome-like structures observed in E3.25 embryos (Figure 2C) show that ICM cells isolated from contact-free surfaces can generate luminal precursor-like structures (Taniguchi et al., 2015 and 2017), which is in line with the observation that all cells of the embryo contribute to lumen formation. Furthermore, the morphology and secretion dynamics of the vesicles reported here are markedly similar to those present in systems that require regulated secretion of contents from large vesicles into a lumen (Miklavc et al., 2012; Rousso et al., 2016; Segal et al., 2018; Tran et al., 2015). If molecules such as morphogens or chemokines are secreted through this mechanism there could be a causal link between fluid accumulation and fate specification and/or cell migration (Durdu et al., 2014). In addition, if vesicles contain highly concentrated osmolytes or cell-adhesion modifying molecules (e.g. podocalyxin) secretion would facilitate downstream lumen expansion as seen in other systems (Takeda et al., 2000; Strilić et al., 2010).

In addition to signaling molecules present in the fluid, the constantly changing physical microenvironment within the blastocyst on account of lumen emergence, coalescence and expansion may also impact tissue remodeling through changes in cell adhesion and cell shape (Durdu et al., 2014; Engler et al., 2006; Mammoto et al., 2013). It has been shown that in epithelial systems multi-luminal phenotypes due to incomplete coalescence can alter the ratios of cell types within tissues and disrupt tissue function (Bagnat et al, 2007; Bryant et al., 2010; Chou et al., 2016; Kesavan et al., 2009). Our recent work (Chan et al., 2019) shows that lumen expansion plays an important role in controlling embryo size and spatial allocation of TE and ICM cells. This combined with the data presented here provide evidence that lumen expansion not only facilitates the molecular specification of PrE cells, but also behaves as a guide for tissue patterning within the embryo.

It would be interesting to consider the possibility that lumen provides both biochemical (e.g. signaling or nutrients) and physical (e.g. contact-free surfaces or pressure) cues sensed by ICM cells, which guide the self-organization of the blastocyst pattern. Further studies to profile the luminal contents and dissect the interconnections of tissue mechanics will need to be conducted in order to understand the precise regulation of luminal coalescence and its influence on cell fate specification.

## Acknowledgements

We thank the EMBL Advanced Light Microscopy Facility and Laboratory Animal Resources Facilities. We thank Jonas Hartmann for discussions regarding analysis methods, assistance in developing the automatic lumen segmentation and providing aliquots of Gateway plasmids for cloning. We thank Dimitri Fabrèges for discussions regarding construction of analysis pipelines. We thank Aissam Ikmi, Jonas Hartmann and members of the Hiiragi Group for critical reading. A.Q.R. was supported by a Boehringer Ingelheim Fonds PhD Fellowship during part of this work (2016-2018). C.J.C. is supported by EMBL Interdisciplinary Postdocs (EIPOD) Fellowship under Marie Sklodowska-Curie Actions COFUND (grant number 664726). The Hiiragi laboratory is supported by EMBL, German Research Foundation, and the European Research Council (ERC Advanced Grant “Selforganising Embryo”, grant agreement 742732).

## Author contributions

Project conceptualization and design, A.Q.R., C.J.C. and T.H. Experiments, A.Q.R. Data analysis, quantification and statistical analysis, A.Q.R. and F.G. Writing, A.Q.R., C.J.C., F.G and T.H. Data interpretation, A.Q.R., C.J.C., F.G., and T.H. Supervision, T.H. and F.G.

---

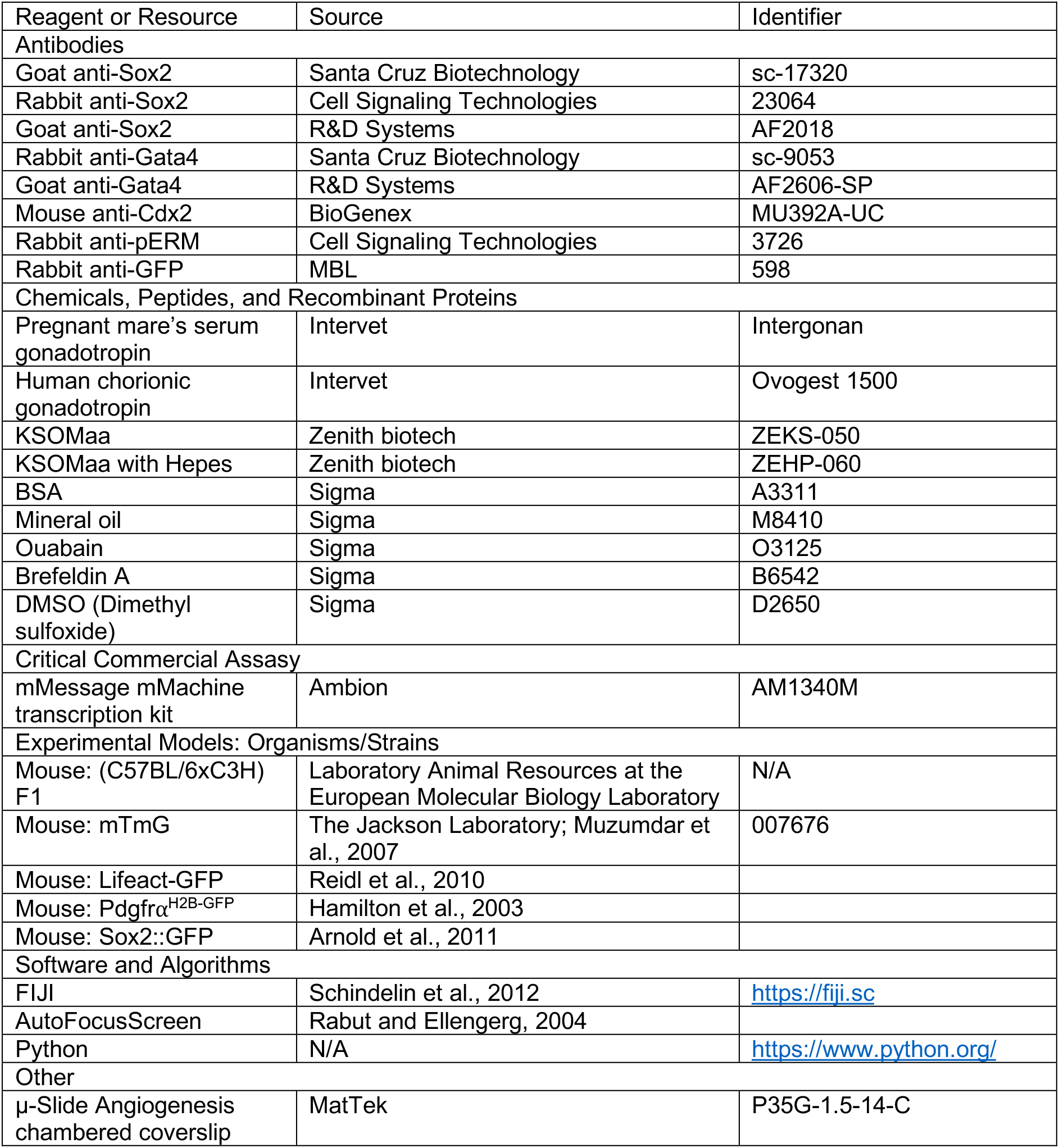
Methods.

## STAR⋆METHODS

### CONTACT FOR REAGENT AND RESOURCE SHARING

Further information and requests for resources and reagents should be directed to the Lead Contact Takashi Hiiragi.

## EXPERIMENTAL MODEL AND SUBJECT DETAILS

### Animal Work

All animal work was performed in the Laboratory Animal Resources (LAR) Facility at European Molecular Biology Laboratory (EMBL) with permission from the institutional veterinarian (ARC number TH110011). LAR Facilities operate according to international animal welfare guidelines (Federation for Laboratory Animal Science Associations guidelines and recommendations). All experimental mice were maintained in specific pathogen-free conditions on a 12-12-hour light-dark cycle and used from 8 weeks of age.

### Transgenic Mice and Genotyping

The following mouse lines were used in this study: (C57BL/6xC3H) F1 as WT, mTmG (Muzumdar et al., 2007), Lifeact-EGFP (Reidel et al., 2010), Pdgfrα^H2B-GFP^ (Hamilton et al., 2003) and Sox2-GFP (Arnold et al., 2011). Standard tail genotyping procedures were used to genotype mice (for primers and PCR product sizes see Table S1).

### Mouse Embryo Recovery and Culture

To obtain pre-implantation embryos, female mice were superovulated by intraperitoneal injection of 5 or 7.5 international units (IU) of pregnant mare’s serum gonadotropin (Intervet, Intergonan) followed by 5 or 7.5 IU of human chorionic gonadotropin (hCG; Intervet, Ovogest 1500) 48 hours later. Hormone injection dosage was batch dependent and determined by LAR services. Superovulated females were mated with male mice directly after hCG injection. Embryos were flushed from dissected oviducts and uteri of female mice after super-ovulation and mating with male mice. Embryos were flushed from dissected oviducts and uteri at 48, 72, 78, 84, 96 and 108 hours post-hCG using KSOMaa with Hepes (Zenith biotech, ZEHP-060). After flushing, embryos were washed in KSOMaa with Hepes, transferred to 10μL drops of KSOMaa (Zenith biotech, ZEHP-050) covered with mineral oil (Sigma, M8410) on either a tissue culture dish (Falcon, 353001) or petri dish (Falcon, 351008) and then cultured at 37°C in a CO2 incubator (Thermo Scientific, Heracell 240i) with 5% CO2.

## METHOD DETAILS

### Pharmacological Inhibition

Ouabain (Sigma, O3125) was resuspended in DMSO (Sigma, D2650) at a stock concentration of 100mM. For working concentrations of 500μM, 250μM and 100μM the stock concentration was diluted in KSOMaa. Brefeldin A (Sigma, B6542) was resuspended in DMSO at a stock concentration of 1.4M. For working concentrations of 14μM and 7μM the stock concentration was diluted in KSOMaa. Embryos were incubated with the appropriate working concentration of ouabain or an equivalent DMSO concentration in μ-Slide chambered coverslips (Ibidi, 81506) for a 12-hour period before fixation in 4% PFA (see Immunofluorescence Staining).

### Serial Mechanical Deflation

Embryos were mounted on epifluorescence microscope (Zeiss, Observer.Z1) equipped with temperature-controlled incubation chamber and visualized using transmitted light. A micromanipulator (Narishige, MON202-D) with a glass holding needle (Harvard Apparatus, GC100T-15) was used to stabilize the embryo while a fine-tipped glass needle attached to a second micromanipulator was used to actively deflate the lumen by penetrating the mural TE at the junction of two cells and manually applying negative pressure. This action was repeated multiple times during the experimental window.

As a procedural control, the deflation needle was inserted at the junction of two mural TE cells in the same manner as experimental embryos and subsequently removed while maintaining a net zero difference in pressure between the lumen and the deflation needle throughout the entire duration of the procedure.

### Cloning and *in vitro* Transcription

The CDS of *fgf4* without a stop codon was cloned into a Gateway middle entry clone and then used in an LR reaction with a 5’ entry clone containing an SP6 site and a 3’ entry clone containing the CDS of *mNeonGreen* with a polyA site.

### mRNA Injection

Linearized plasmid was used as the template for an *in vitro* transcription reaction from an Invitrogen mMessage SP6 kit (AM1340M). mRNA injections were performed on the same microscope and with the same micromanipulators as described earlier (see Serial Mechanical Deflation). mRNA was injected into a single blastomere of 4-cell stage embryos at a concentration of 200ng/μl using a needle (Harvard Apparatus, G100TF-15) attached to an injector (Eppendorf, FemtoJet).

### Immunofluorescence Staining

Embryos were fixed with 4% PFA for 15 minutes at room temperature and subsequently permeabilized with PBS (0.5% Triton-X) for 20 minutes at room temperature before transferring to blocking buffer (PBS with 0.1% Tween-20; 5% BSA) for at least 4hrs at 4°C. Embryos were incubated with primary antibodies diluted in blocking buffer overnight at 4°C. After washing with blocking buffer, embryos were incubated with secondary antibodies diluted in blocking buffer for 2hrs at room temperature. Finally, embryos were rinsed with PBS before being mounted in a DAPI solution (PBS with 1:2000 DAPI; Invitrogen, D3571) for imaging (see Fixed Sample Imaging).

The following primary antibodies were used in this study: rabbit anti-pERM (Cell Signaling, 3726), mouse anti-Cdx2 (BioGenex, MU392A-UC), goat anti-Sox2 (Santa Cruz Biotechnology, sc-17320), goat anti-Sox2 (R&D Systems, AF2018-SP), rabbit anti-Sox2 (Cell Signaling, 23064), rabbit anti Gata4 (Santa Cruz Biotechnology, sc-9053), goat anti-Gata4 (R&D Systems, AF2606-SP), rabbit anti-GFP (MBL, 598). Secondary: donkey anti-goat Alexa Fluor 488 (Life Technologies, A-11055), donkey anti-rabbit Alexa Fluor 488 (Life Technologies, R37118), donkey anti-rabbit Alexa Fluor 546 (Life Technologies, A10040) and donkey anti-mouse Cy^TM^5 AffiniPure (Jackson Immunoresearch, 715-175-150). All secondary antibodies were used at 1:200 dilutions. DAPI was used to visualize nuclei. Rhodamine phalloidin (Invitrogen, R415) was used to visualize F-actin at a 1:200 dilution.

### Microscopy

#### Fixed Sample Imaging

Either point scanning confocal imaging on an LSM-780 (Zeiss) or Airyscan Fast Mode acquisition on an LSM-880 (Zeiss) was performed for immunofluorescence staining prepared samples.

#### Live Imaging

Embryos were mounted in either 10μL KSOMaa drops covered with mineral oil (Sigma, M8140) on 35mm glass-bottomed dishes (MatTek, P35G-1.5-14-C), or in 15-well chambered coverslips (Ibidi, 81506) with 60μL of KSOM if part of a pharmacological inhibition experiment. For long-term imaging (timesteps >5 minutes), imaging was performed on an LSM-780 (Zeiss) with XY sample drift compensation (Rabut and Ellenberg, 2004). Short-term time-lapse imaging (timestep = 10 seconds) was performed on an lSm-880 using Airyscan Fast Mode acquisition. Both microscopes are equipped with custom-made temperature and gas-controlled chambers (EMBL) set to 37°C and 5% CO2 during all experiments and C-Apochromat 40x water objectives (Zeiss).

## QUANTIFICATION AND STATISTICAL ANALYSIS

### Image Analysis

#### Volume Quantification

Volume quantification for time lapses was calculated from the output of a custom-written automatic membrane segmentation pipeline (Table S2). Volume threshold for segmentation was set to 1pL, 5pL, or 10pL depending on the image quality of the dataset to ensure fidelity.

For vesicles and single timepoints volume was estimated manually by measuring the vesicular or luminal circumference of the central plane in FIJI to extract the radius assuming isotropy.

#### Polarity Phenotype Scoring

Immunofluorescence images were examined for microlumina, apicosome-like structures and lumen-facing membranes expressing pERM. Binary scores were assigned to images on a presence (1), absence (0) basis for each structure, and the frequency of occurrence determined from the binary scores.

#### Center of Mass Distance to Lumen Surface

The lumen was segmented as described in *Volume Quantification.* The ‘segmented cell mass’ is taken to be the sum of all cells in the embryo such that the total embryo volume is equivalent to the sum of the ‘segmented cell mass’ and the ‘segmented lumen.’ ‘Lumen boundaries’ were determined from the lumen segmentation. ‘Cell mass boundaries’ were determined from the sum of the cell mass and lumen segmentations. Convex hulls representing the lumen and the embryo surface were created from the two boundary point sets. The center of mass of the lumen (*P*_1_) and the center of mass of the cell mass (*P*_2_) were used to determine the embryonic-abembryonic axis 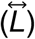. Lumen boundary and embryo outer boundary intersection points with 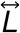 (*P*_3_ and *P*_4_ respectively) were found by recursively searching the facets of the convex hulls such that 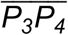 contains *P*_2_ but does not contain *P*_1_. The ICM signal of interest was automatically segmented and the weighted center of mass (*P*_5_) calculated. 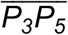 is taken as the absolute distance of the signal of interest to the ICM-lumen interface, which is then normalized by the absolute ICM width 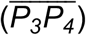. This normalization is done to ensure no intrinsic bias due to changes in tissue morphology or size are introduced. The normalized distance for timepoint *t_n_* in a time-lapse starting at t_0_, is 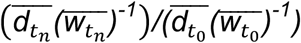 where 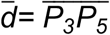 and 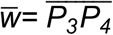. All segmentations and calculations were performed automatically using custom Python scripts (Table S2).

As the ICM width along the embryonic-abembryonic axis inherently shrinks throughout blastocyst development, the lack of global movement of all tracked signals toward the lumen confirms ICM width normalization to be a valid measure for positional normalization when examining signals within ICM populations relative to other objects along or relative to the embryonic-abembryonic axis.

#### Fate Specification Analysis

For immunofluorescence images, nuclei were segmented using DAPI signal as a reference. Transcription factor signal was then measured in each nucleus for all channels. From these measurements, a sum measurement was calculated for each channel and subsequently normalized to the reference DMSO control category. The entire analysis was performed using custom Python scripts (Table S2).

#### Spatial Segregation Analysis

Using the nuclear segmentations acquired during *Fate Specification Analysis,* the center of mass was calculated for each nucleus and stored as a 3D coordinate. The median and mean of each dimension from all nuclear centers of mass were calculated resulting in two 3D points of reference within the embryo, such that the median (P”) will be within the lumen and the mean (*P_2_*) will be within the ICM. A 3D line representing the embryonic-abembryonic axis 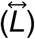 was determined from *P*_1_ and *P*_2_. A 3D line segment 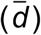 is drawn from the center of mass of a nucleus (*P*_3_) to an intersection point with 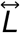 (*P*_4_) such that the angle of intersection is 90°. The 3D embryo can then be represented on a 2D coordinate system in which the *x* axis is 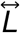 such that *x* = 0 is the minimum of all *P*_4_ points identified and *y* values are the length of 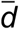. After this 3D to 2D projection, the fluorescence intensity values of channels of interest were plotted along a third axis *(z)* against the positional information (*x,y*). A multidimensional kernel density estimate (KDE) is then performed to acquire a probability density map. KDEs from different channels of interest are then integrated over one another in 3D to obtain a single scalar value representing the degree of sorting between groups (‘Overlap’). Maximum potential overlap values (‘Simulation’) are derived from the ranges of expression and domain sizes observed in WT embryos. The entire analysis was performed using custom Python scripts (Table S2).

#### Image Processing for Figures

Images for Figures 1A, 1B and 3 were processed with a multidimensional median filter (size = 10, scipy.ndimage). Images for Figures 1C, 1F and 2 were acquired using Zeiss Airyscan Fast Mode and processed using the corresponding automatic 3D deconvolution.

### Cell Counts

Total embryo cell numbers for Figures S2,S3 and S4 were determined from the automatic segmentation of nuclear masks (Table S2) based on DAPI signal. The results from automatic segmentations were validated manually in FIJI for a subset of images in each experimental category to check that the difference between the methods was negligble.

### Statistical Analysis

All graphs were generated and statistical analysis was performed in Python using the scipy statistics package. Two-sided Fisher exact test was used to compare the frequency of observation between two binary datasets. Kruskal-Wallis H-test for independent samples (non-parametric ANOVA) was used to test for statistical significance between populations; ****p<0.0001 ***p<0.001 **p<0.01 *p<0.05. Sample sizes and p-values are indicated in text, figures and figure legends. No statistical tests were used to predetermine sample sizes.

---

**Figure S1.**
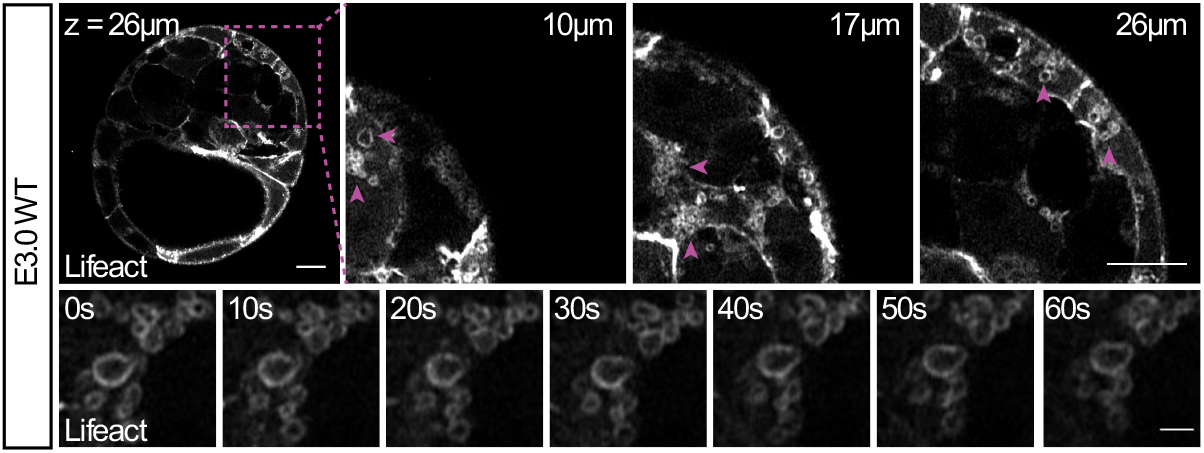
Cortically localized and secreted vesicles are no longer present in embryos containing an expanded lumen. Z-slices of an E3.5 WT embryo expressing Lifeact-GFP show only cytoplasmic vesicle clusters (top row, scale bars = 10μm.). Inset (panels 2-4) delineated by magenta box (panel 1). Arrowheads (panels 2-4) highlight vesicle clusters. Time-lapse of vesicle cluster dynamics in an E3.5 WT embryo expressing Lifeact-GFP (bottom row, scale bar = 2μm).

**Figure S2.**
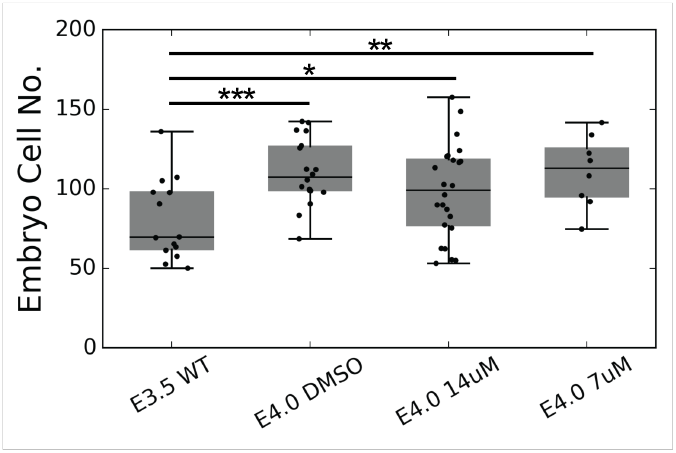
COPII inhibition via Brefeldin A treatment does not affect embryo cell number. Total embryo cell counts for E3.5-E4.0 COPII inhibition conditions – E3.5 WT, N = 14; E4.0 DMSO, N = 18; E4.0 7μM, N = 8; E4.0 14μM, N = 24. *p<0.1 **p<0.01 ***p<0.001

**Figure S3.**
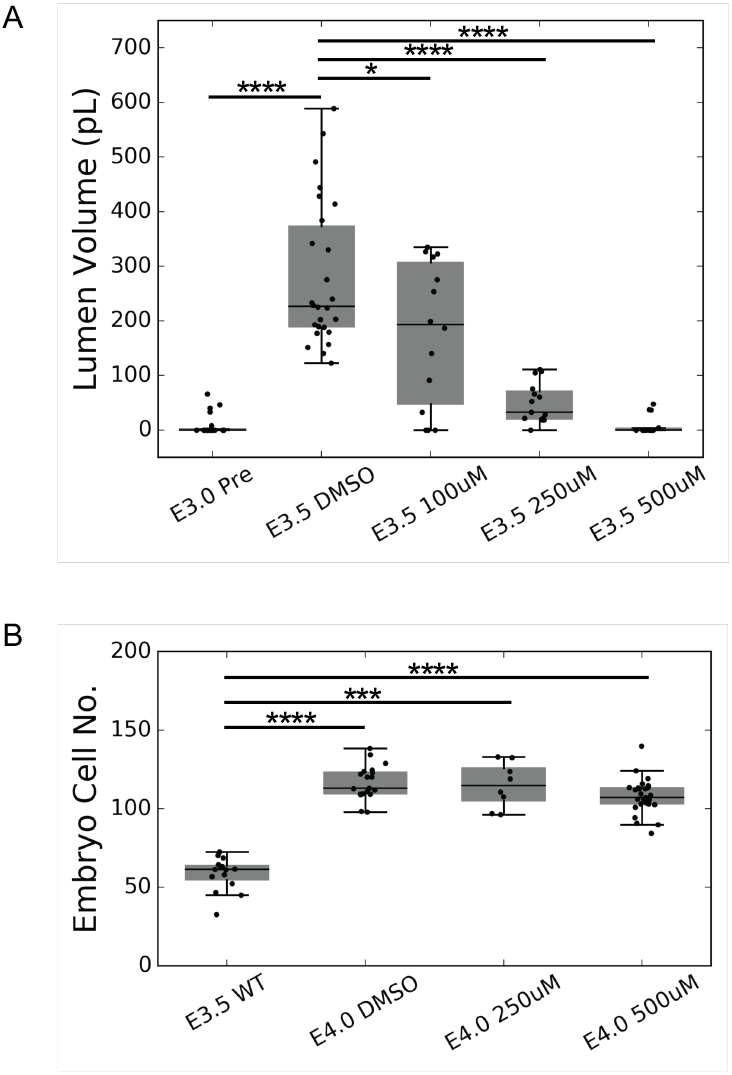
Inhibitory effects of oubain are titratable by dose and do not affect cell number. (A) Volume of lumina for experimental and control embryos in E3.0-E3.5 ouabain titration inhibitions – E3.0 WT, N = 20; E3.5 DMSO, N = 26; E3.5 100μM, N = 14; E3.5 250μM, N = 15; E3.5 500μM, N = 14. (B) Total cell numbers of embryos in experimental and control groups of ouabain titration inhibitions – E3.5 WT, N = 15; E4.0 DMSO, N = 19; E4.0 250μM, N = 8; E4.0 500μM, N = 27. *p<0.05 ***p<0.001 ****p<0.0001

**Figure S4.**
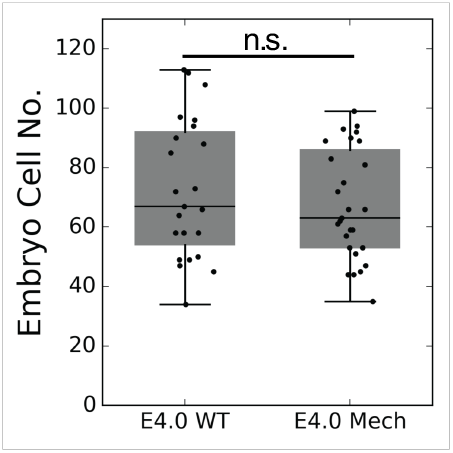
Serial mechanical deflation does not impact cell number. Boxplot of total cell numbers of embryos in experimental and control groups for serial mechanical deflation – E4.0 WT, N = 23; E4.0 Mech, N = 27. n.s. = not significant.

## Supplementary tables

**Table S1.**
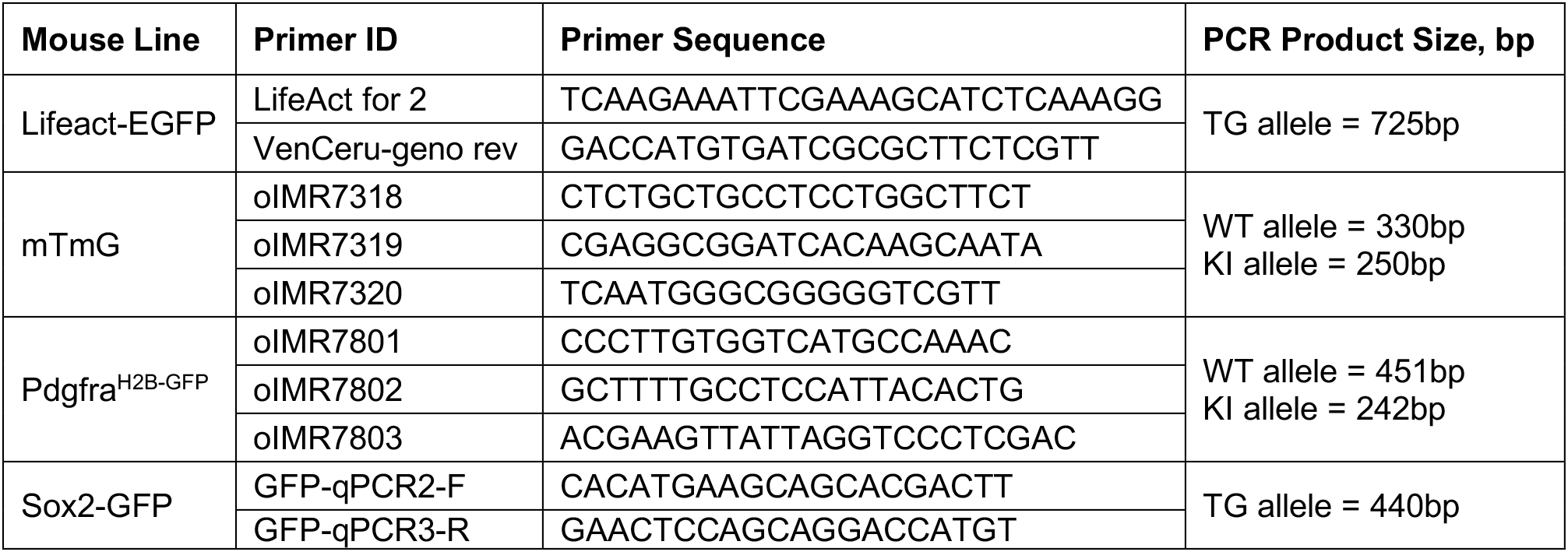
Genotyping primers, related to STAR Methods.

**Table S2.** Key Python 2.7 functions and packages used in custom image analysis pipelines.

| Analysis | Package | Key Functions |
| --- | --- | --- |
| Lumen Segmentation | scipy.ndimage | median_filter, binary_opening, binary_closing, label |
|  | numpy | percentile, sum |
| Reporter Expression Center of Mass | scipy.ndimage | median_filter, measurements.center_of_mass |
| Centroid Distance to Lumen Surface | scipy.spatial | distance.cdist, ConvexHull |
|  | skimage.segmentation | find_boundaries |
|  | numpy | where |
|  | scipy.ndimage | binary_fill_holes |
|  | sympy | geometry.line3d.Line3D, Point3D, intersection |
| Fate Specification | scipy.ndimage | distance_transform_edt, label |
|  | skimage.feature | peak_local_max |
|  | skimage.morphology | watershed |
|  | numpy | unique, mean, sum |
| Spatial Segregation | scipy.ndimage | measurements.center_of_mass |
|  | numpy | mean, median |
|  | sympy.geometry | line3d.Line3D, distance, perpendicular_segment, intersection |
| Luminal Proximity | scipy.ndimage | binary_dilation |
| Cell Counts | scipy.ndimage | median_filter, binary_opening, measurements.label |

## Supplementary movies

**Movie S1. Vesicles localize to all basolateral and apolar membranes prior to visible extracellular fluid accumulation.**

Z-stack of actin signal (phalloidin) in a WT E3.0 embryo. Scale bar = 10μm.

**Movie S2. Vesicles are actively secreted into intercellular space.**

Time-lapse (timestep = 10 seconds) of actin localization (Lifeact-GFP) during vesicle secretion in a WT E3.0 embryo. Scale bar = 5μm.

**Movie S3. WT localization of vesicles is maintained in Atp1 inhibited embryos.**

Z-stack of actin signal (Lifeact-GFP) in an Atp1 inhibited E3.0 embryo. Scale bar = 10μm.

**Movie S4. Vesicles continue to be secreted into intercellular space during Atp1 inhibition.**

Time-lapse (timestep = 10 seconds) of actin localization (Lifeact-GFP) during vesicle secretion in an Atp1 inhibited E3.0 embryo. Scale bar = 5μm.

**Movie S5. Apicosome-like structures are contained in cells isolated from luminal contact.**

Z-stack of a cell containing an apicosome-like structure (pERM, magenta; Actin, gray; nuclei, cyan). Scale bar = 5μm.

**Movie S6. Apicosome-like structures are released into luminal space when the cell gains a contact-free surface.**

Time-lapse (timestep = 15 minutes, hh:mm) of membrane signal (mT) in a cell releasing an apicosome-like structure into luminal space once the cell acquires a contact-free surface along the ICM-lumen interface. Scale bar = 10 μm.

